# Evaluating transformer-based models for structural characterization of orphan proteins

**DOI:** 10.64898/2026.03.10.709490

**Authors:** Ercan Seçkin, Dominique Colinet, Etienne GJ Danchin, Edoardo Sarti

## Abstract

**Motivation:** Transformer-based models (TBMs) are state-of-the-art deep learning architectures that predict protein structural features with high accuracy. Despite methodological differences, they all rely on large protein sequence datasets structured by homology, as homologous proteins typically share similar structures. However, 5–30% of eukaryotic proteomes consist of orphan proteins—sequences without detectable similarity to known families. Although they may share structural traits with characterized proteins, their lack of homology makes them and ideal dataset for evaluating TBM generalization beyond familiar sequence space.

**Results:** We compared predictions from several widely used TBM architectures on an expert-curated set of orphan proteins from the Meloidogyne genus. None of these proteins has an experimentally determined structure. To assess model performance, we conducted consistency analyses, comparing predicted features with those observed in sets of known homologous proteins and across models. Multiple sequence alignment–based approaches such as AlphaFold2 performed poorly on orphan proteins, as did single-sequence or embedding-based language models including ESMFold, OmegaFold, and ProtT5. This limited performance cannot be fully attributed to intrinsic disorder, as confirmed by independent non-TBM disorder predictors. While accurate tertiary structure prediction remains out of reach, secondary structure is more reliably captured: predictors share about 70% of secondary structure elements on average, regardless of global fold similarity, and these elements are consistently identified by dedicated secondary structure tools.

**Availability:** All data and analysis scripts are available at https://doi.org/10.5281/zenodo.18788931

## Introduction

Since the introduction of the transformer architecture, protein structure prediction has undergone a major conceptual and practical leap. Transformer-based models (TBMs), originally developed for natural language processing, have proven particularly effective at capturing long-range dependencies in protein sequences and at integrating evolutionary and structural information at scale (Akdel et al., 2022). Their adoption has led to a new generation of predictors that significantly outperform previous approaches based on coevolutionary statistics and shallow neural architectures.

A milestone in this transition is AlphaFold2 (Jumper et al., 2021), which introduced a set of ad-hoc architectural components tailored to protein geometry. AlphaFold2 jointly reasons over sequence and pairwise representations using attention mechanisms, including pairwise and triangular attention, and incorporates an equivariant structure module based on invariant point attention (IPA) and explicit SE(3) frames. This design allows the model to iteratively refine atomic coordinates while preserving rotational and translational equivariance, resulting in near-experimental accuracy for many targets.

Subsequent models have explored alternative ways of leveraging transformer architectures. ESMFold (Lin et al., 2023) builds upon large protein language models such as ESM-1b and ESM2 to generate sequence and pairwise embeddings directly from single sequences, without relying on multiple sequence alignments. These embeddings are then processed by a structure module derived from AlphaFold2, together with a newly introduced folding block. For confidence estimation, ESMFold reports pLDDT values computed using AlphaFold2’s prediction head applied to its own internal representations.

A different design philosophy is exemplified by OmegaFold (Wu et al., 2022), which employs a transformer-based architecture trained end-to-end for structure prediction while explicitly incorporating geometric constraints and rotation-equivariant operations. OmegaFold emphasizes rapid inference and reduced dependence on evolutionary depth, further highlighting the versatility of transformer architectures for structural modeling.

Beyond full 3D structure prediction, transformer-derived protein language models have been widely used to extract informative embeddings for downstream tasks. Models such as ProtT5 (Elnaggar et al., 2022), trained on hundreds of millions of sequences, have demonstrated strong performance in predicting secondary structure, transmembrane regions, and other local structural features, even in low-similarity regimes.

Despite these advances, TBM-based predictors are not free from limitations. Prediction biases and structural hallucinations have been repeatedly observed, particularly for proteins with shallow evolutionary context or unusual folds (Pratt et al., 2025). While recent studies have begun to quantify the extent of these errors and to investigate their underlying causes (Williams et al., 2025) (Sarti and Cazals, 2026) (Cazals and Sarti, 2025), a comprehensive theoretical explanation of why and when such failures occur is still lacking.

In parallel, progress in protein structure analysis has been enabled by new methods for fast and sensitive structural comparison. FoldSeek (van Kempen et al., 2024) introduced the 3Di structural alphabet, which encodes local backbone conformations as discrete symbols, allowing structural alignments to be reduced to sequence-like comparisons. Other approaches similarly exploit compact structural representations to improve scalability and classification accuracy. Here too, transformer-based models play a role: ProstT5 (Heinzinger et al., 2024) predicts 3Di sequences directly from amino acid sequences and vice versa, bridging sequence- and structure-based representations within a unified framework.

All TBMs ultimately rely on the abundance of available protein sequence data for training, yet their architectures are inherently sensitive to out-of-distribution inputs. Orphan proteins, by definition, lack detectable homologs in current databases and are therefore absent from training sets; moreover, no experimentally determined structures are available for this class of proteins (Fakhar et al., 2023). Orphan proteins can arise through two distinct etiologies. Some correspond to highly diverged members of known protein families, whose evolutionary origins are obscured by strong selective pressures and extensive sequence divergence, while others are *de novo* proteins that have emerged relatively recently from previously non-coding genomic regions (Seçkin et al., 2025b). In the former case, it is generally assumed that aspects of the ancestral fold and function may be conserved despite the loss of detectable sequence similarity, whereas for *de novo* proteins there is little prior knowledge regarding either structure or function.

These two origins also suggest different expectations for intrinsic disorder. For diverged proteins, the overall level of intrinsic disorder is expected to be broadly comparable to that of their (unknown) ancestral family. In contrast, several studies have reported that *de novo* proteins are enriched in intrinsically disordered regions, consistent with their amino acid composition and recent evolutionary origin (Wilson et al., 2017) (Heames et al., 2020) (Basile et al., 2017). However, this view is not unanimous, and other analyses have found no systematic increase in disorder relative to older proteins (Ekman and Elofsson, 2010) (Schmitz and Bornberg-Bauer, 2018). Taken together, these considerations make orphan proteins an ideal dataset for assessing the true generalization capabilities of transformer-based approaches. In this work, we compare several TBM-based predictors on a recently published orphan protein dataset. While tertiary structure predictions remain unreliable in this regime, we find that secondary structure can still be captured with notable accuracy.

## Methods

### Meloidogyne orphan proteome benchmark

Meloidogyne protein datasets and their orphan status were obtained from a previous comparative genomics study in which eight Meloidogyne species were analyzed against all other available nematode proteomes and a broad representation of non-nematode taxa using a dedicated homology-detection pipeline (Seçkin et al., 2025a). In that study, orphan proteins were defined as those with no detectable homologs outside the genus Meloidogyne, as determined by sequence similarity searches and orthology inference. Homologous proteins were clustered into orthogroups and a total of 8,974 orthogroups were identified as orphans, which corresponds to a total of 48,681 proteins, of which approximately 20% were inferred to result from extreme sequence divergence, while 18% were suggested to have emerged *de novo*. For the rest of the orphans, it was difficult to identify the origin and emergence as it is not always evident how to trace them back to the distant homolog or non-coding region from which they appeared. For the control benchmark of non-orphan proteins, we only considered sequences that were present in the Meloidogyne genus and had detectable homologs in at least four additional non-Meloidogyne species.

### Structure predictions

Three independent structure prediction approaches were used to model orphan and non-orphan proteins: AlphaFold2 (Jumper et al., 2021), OmegaFold (Wu et al., 2022), and ESMFold (Lin et al., 2023).

For AlphaFold2 (v 2.3.2), structure prediction was performed at the orthogroup level rather than per individual sequence. For each orthogroup, a custom multiple sequence alignment (MSA) was generated using the protein sequences belonging to that orthogroup with MAFFT (Katoh et al., 2017). One representative sequence per orthogroup was randomly selected as the query sequence, and the corresponding orthogroup-specific MSA was used as input to AlphaFold2. This strategy allowed us to leverage evolutionary information within orthogroups in the absence of homologs outside of the genus. Other parameters were used on default.

OmegaFold (v 1.1.0) and ESMFold (v 1.0) predictions were performed independently for each protein sequence without the need of using custom MSAs, following the default parameters of each method. For orphan proteins, one structural model was therefore generated per protein sequence. In order to compose a comparable negative sample of non-orphan structure predictions, we created two different sets:

- Same-length Meloidogyne sequences. For each representative sequence of the 8,974 orphan orthogroups, we selected one non-orphan sequence from the 8 available Meloidogyne species having approximately the same length. We use this set whenever the important difference orphan/non-orphan in sequence length can affect analyses;
- *M. incognita* whole non-orphan proteome. *M. incognita* is the species with the most comprehensive genomic and functional annotation. For this dataset, one representative sequence per orthogroup was selected, and structure prediction was performed using AlphaFold2, OmegaFold, and ESMFold under the same settings described for orphan proteins.

Structural confidence was evaluated using the mean predicted Local Distance Difference Test (pLDDT) score for each model, as provided by the respective tools. Structural similarity between predictions obtained from different methods was assessed using TM-score calculated with TM-align (release 2018-04-26) (Zhang and Skolnick, 2005).

### Disorder predictions

Intrinsic disorder was predicted from amino acid sequences using three independent predictors: flDPnn (Hu et al., 2021), AIUPred (Erdős and Dosztányi, 2024) (v 1.0), and LoRa-DR (Lombardi et al., 2025) (v 1.0). flDPnn and AIUPred were run using default parameters. LoRa-DR predictions were performed using the model trained on ESM embeddings to estimate residue-level disorder probabilities.

All three predictors provide per-residue disorder probability scores ranging from 0 to 1. A sequence-level disorder metric was calculated by averaging residue-level probabilities across each full-length protein. Mean disorder scores were subsequently compared among highly diverged orphan proteins, *de novo* orphan proteins, and non-orphan proteins.

Disorder can also be estimated through the calculation of the relative surface accessibility (RSA). We use the algorithm described in (Piovesan et al., 2022) on ESMFold predictions as a further comparison.

### Secondary structure predictions

Secondary structure of orphan and non-orphan proteins was predicted directly from sequence using ProtT5 with option to distinguish the three states: helix, sheet and coil. ProtT5 (Elnaggar et al., 2022) generated an output on a sequence format with these three states. The predicted secondary structures from the sequence were then compared to secondary structure predictions of AlphaFold2 and ESMFold using DSSP (Hekkelman et al., 2025), in order to assess the similarity of the predictions.

### Search against PDB and AFDB databases

To assess potential tertiary structure similarities between orphan proteins and previously characterized proteins, we performed structural homology searches using two complementary approaches. In the first approach, orphan protein sequences were converted into 3Di structural representations using ProstT5 (Heinzinger et al., 2024). The resulting 3Di sequences were queried against the Protein Data Bank (PDB) and AlphaFold2 Protein Structure Database (AFDB) using Foldseek (van Kempen et al., 2024) (easy-search mode). Foldseek outputs included alignment statistics such as percent sequence identity and alignment scores, which were used to evaluate the significance of detected similarities.

In the second approach, structural homology searches were performed directly using the predicted three-dimensional models generated by ESMFold. These predicted structures were queried against the same reference databases using Foldseek.

## Results

### TBM predictions of orphan proteins have low pLDDT regardless of MSA usage

We performed AlphaFold2, ESMFold and OmegaFold predictions on an exhaustive set of orphan proteins of the 4 species composing the clade 1 of the genus Meloidogyne, and on the entire non-orphan proteome of *M. incognita* (see Methods and (Seçkin et al., 2025a)). Orphan protein orthologs within the Meloidogyne genus have been grouped in so-called orthogroups (OGs), allowing us to build a shallow input MSA for AlphaFold2. Notably, AlphaFold2 produces one structure prediction per OG, whereas ESMFold and OmegaFold produce one prediction per sequence. When comparing the results, we randomly choose one structure out of the orthogroup.

For the three models, the median and average pLDDT of diverged and *de novo* predictions are in the “low” (50 *< pLDDT <* 70) or “very low” (*pLDDT <* 50) quality ranges, with ESMFold getting systematically lower quality estimates (Figure 1A). The non-orphan predictions are associated with significantly higher pLDDT values, just above or below the “high” quality threshold (*pLDDT* = 70). Despite differences in their averages, the pLDDT scores relative to the three models are found to correlate (Figure 1B and C).

**Figure 1.**
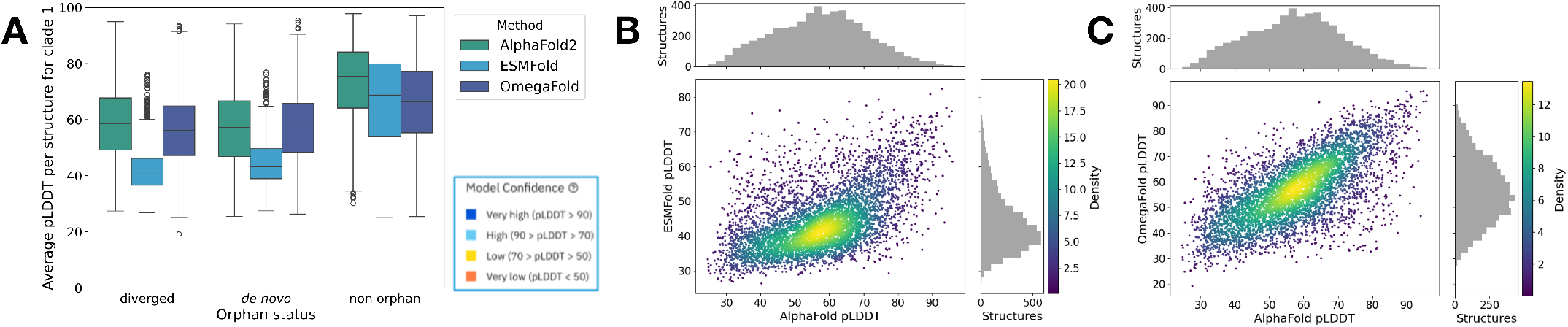
Average pLDDT of orphan protein structure predictions for three TBMs. A. Average pLDDT of AlphaFold2, ESMFold and OmegaFold on confirmed diverged and *de novo* proteins. Average pLDDT of non-orphan proteins for reference. B. Correlation between the orphan protein structure prediction pLDDT of AlphaFold2 and ESMFold (left) and Omegafold (right).

The correlation between OmegaFold and ESMFold reveals two subpopulations (Figure S4). Both show a correlation in pLDDT scores between the methods, but one exhibits a systematic shift in OmegaFold values. This effect arises from larger score differences among members of the same OG in OmegaFold than in ESMFold: the subpopulation with higher OmegaFold values is predominantly composed of structures ranked first within their OGs. This effect is not related to the presence of these sequences in UniProt (and thus in the OmegaFold training set).

### Structural similarity correlates with pLDDT

Given the uniformly low pLDDT scores produced by all three methods, a natural question is whether the pLDDT metric fails to adequately assess structures that may be distant from its training set (the PDB), or whether the predicted structures themselves are of low quality. To investigate this, we performed pairwise structural alignments using TM-Align (Zhang and Skolnick, 2005) between predictions generated for the same protein sequence. We observed a correlation between pLDDT and alignment TM-score: high pLDDT values are associated with strong structural agreement among the three methods, whereas low pLDDT values correspond to divergent predictions (Figure 2). This result supports the reliability of pLDDT and indicates that overall low pLDDT scores reflect poor structural quality even for orphan proteins.

**Figure 2.**
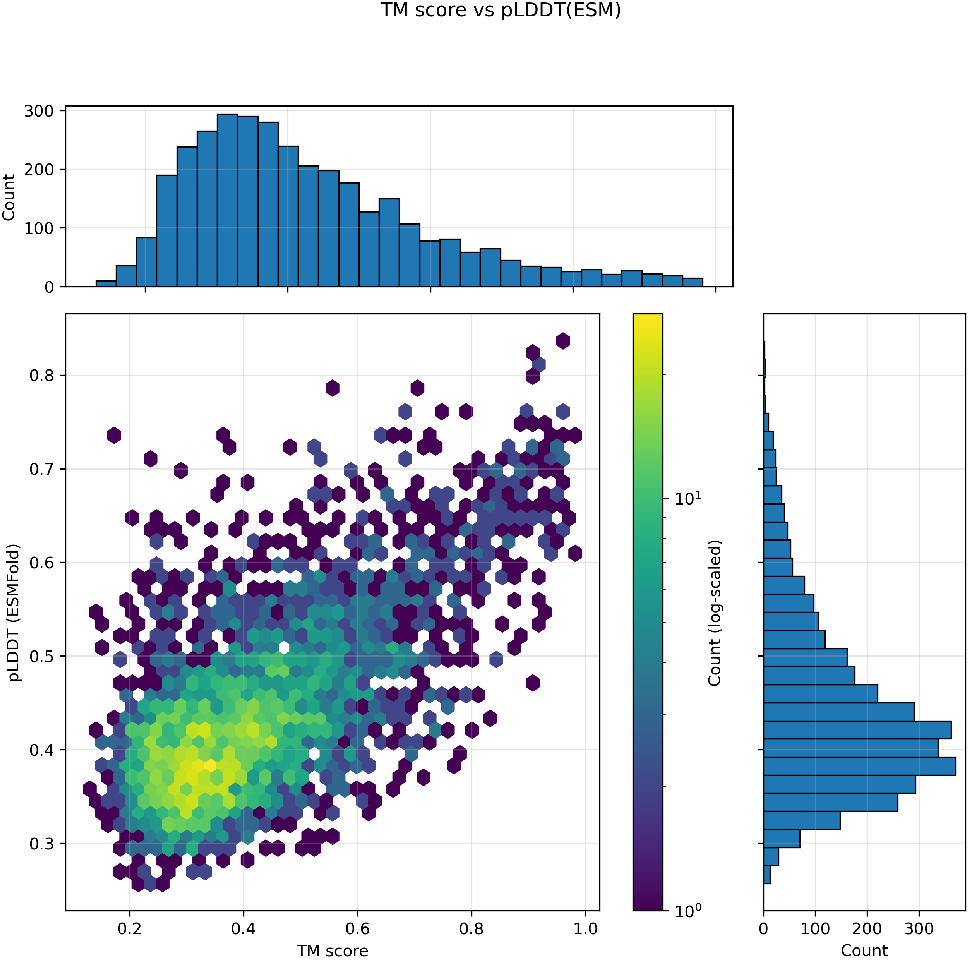
pLDDT correlates with TM-score. TM-score calculated between ESMFold and Omegafold structures plotted against the pLDDT score reported by ESMFold. The more the two predictors output divergent structures, the lower the pLLDT of the predicted structures.

### Intrinsic disorder and orphan structure prediction

Intrinsic disorder (ID) is believed to be prevalent in orphan and especially *de novo* proteins. Lacking direct experimental evidence on Meloidogyne proteins or orphan proteins in general, we used three ID predictors that are progressively less connected to transformer-based structure predictors: LoRA-DR (Lombardi et al., 2025) directly uses PLM embeddings, AIUPred (Erdős and Dosztányi, 2024) uses a transformer architecture trained with data that is not derived from TBMs, and flDPnn (Hu et al., 2021) uses a fully connected encoder on non-TBM data. As a further comparison, we also calculate the relative surface accessibility of the structure predictions. We observe that the more the method is connected to TBMs, the more disorder will be predicted (Figure 3). We also notice that a difference in intrinsic disorder density is only visible for the PLM-based method LoRA-DR and for the RSA calculated directly on the TBMs output, whereas the other two state-of-the-art methods do not find statistically significant differences.

**Figure 3.**
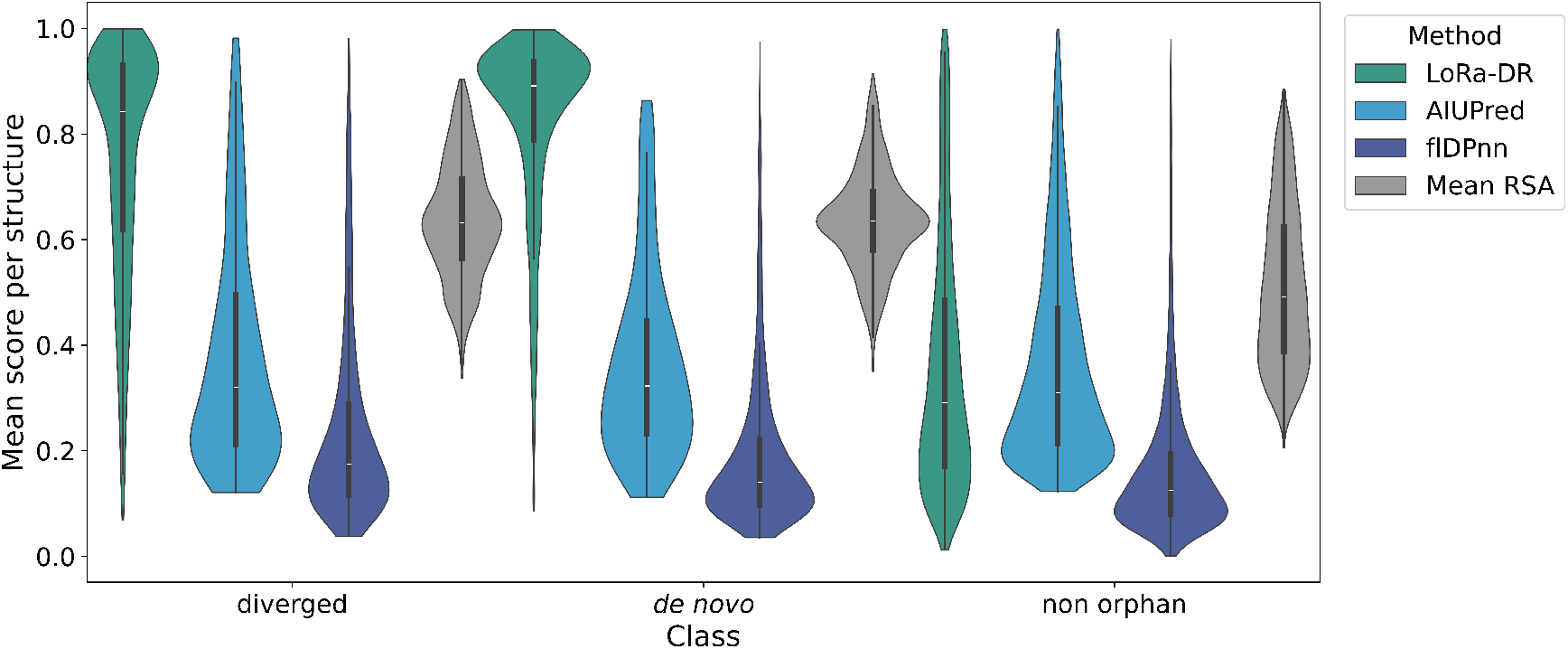
Average predicted intrinsic disorder of protein structure predictions. Predicted intrinsic disorder of Meloidogyne proteins, divided in diverged, *de novo*, and non-orphan. Predicted disorder increases the more the predictive method is closer to TBMs. For the TBM-based method LoRA-DR and the RSA calculation directly on the TBM output structures, orphan proteins are consistently more disordered than non-orphan, whereas for the other methods no difference is detected.

### Predicted orphan tertiary structures lack structural homologs

By definition, orphan proteins lack detectable homologs at the sequence level. Consistent with this, we observed reduced confidence in structure predictions for orphan proteins compared to non-orphan controls. However, structural similarity is often more conserved than primary sequence similarity. We therefore investigated whether orphan proteins might nonetheless share detectable structural similarity with known protein structures.

To address this, we first applied a sequence-based structural search strategy that does not rely on predicted three-dimensional models. Representative sequences from each of the 8,974 orphan orthogroups were converted into 3Di representations using ProstT5. These 3Di sequences were then queried against the Protein Data Bank (PDB) and the AlphaFold Protein Structure Database (AFDB) using Foldseek. This approach yielded only 64 hits against PDB and 79 hits against AFDB across all orphan orthogroups. Among these, only two PDB hits and one AFDB hit exhibited greater than 50% identity, indicating that high-confidence structural matches are extremely rare.

In a second approach, we performed structural searches using predicted ESMFold models for orphan proteins, despite their overall lower prediction confidence. Querying these models against PDB and AFDB resulted in a larger number of matches (2,404 hits against PDB and 3,893 hits against AFDB). However, high-identity matches remained uncommon, with only 23 hits exceeding 50% identity against PDB and 16 against AFDB.

Taken together, these results indicate that the vast majority of orphan proteins lack detectable high-confidence structural homologs in current structural databases. Even when leveraging predicted models, structural similarity to known proteins remains limited. Consequently, reliable structural inference for orphan proteins remains challenging.

### Similar secondary structure content, lower confidence

The predicted structures of orphan proteins do not exhibit a systematic increase in flexible or disordered regions. Specifically, the proportion of residues participating in secondary structure elements (SSEs) remains largely unchanged across the three TBMs considered: AlphaFold2, ESMFold, and ProtT5 (Figure 4). Notably, ProtT5 is the only method that does not overestimate alpha helical SSEs and that minimizes differences in beta sheet SSE content.

**Figure 4.**
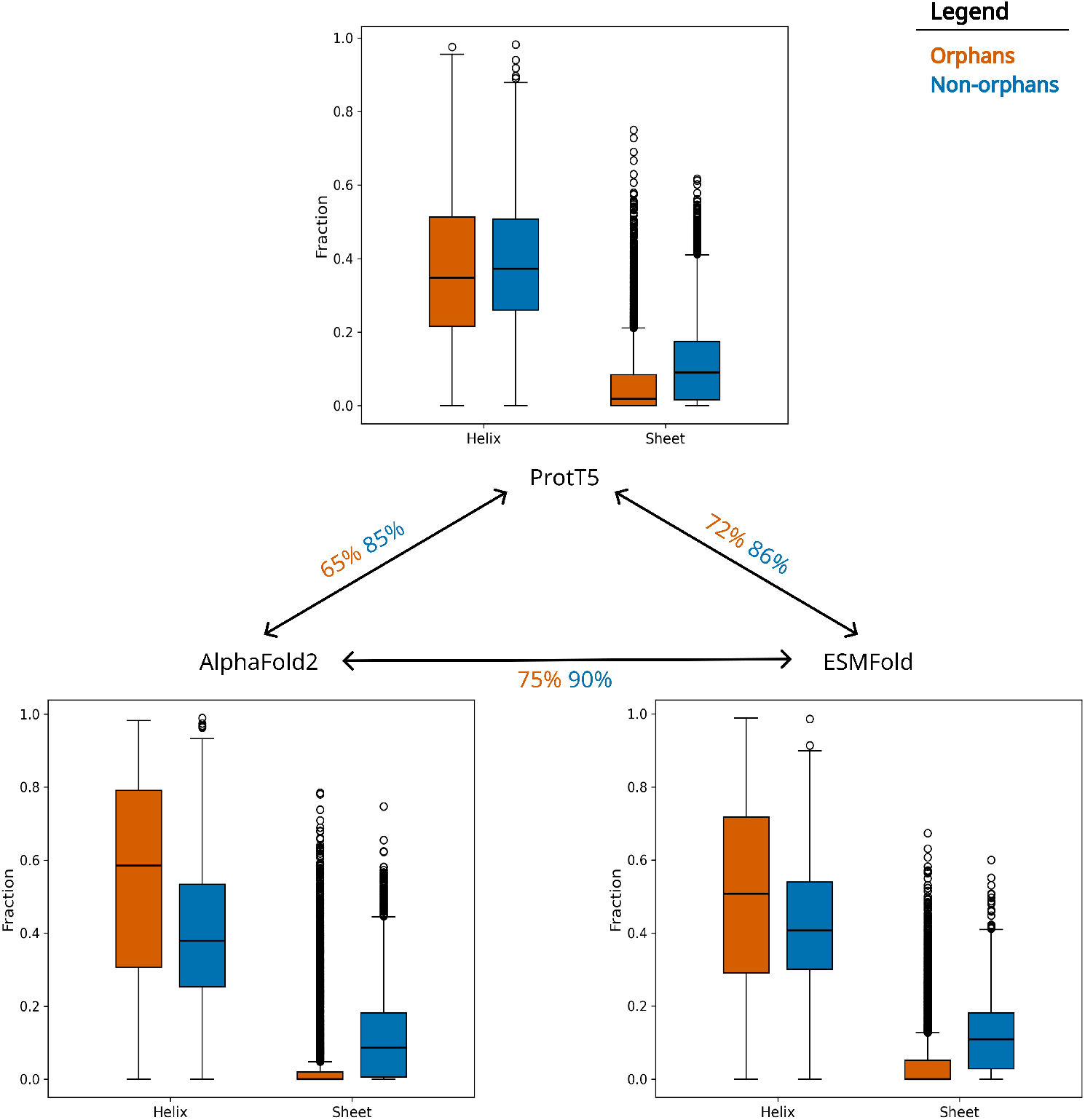
Predicted secondary structure on orphans and non-orphans for three types of TBMs. Fraction of amino acids per sequence that are predicted as alpha helix and beta sheet, for orphan (orange) and non-orphan (blue) proteins. Percentages on connecting arrows describe the average consensus of the two methods on alpha/beta/coil assignment.

For non-orphan proteins, SSE assignments are largely consistent across TBMs, with pairwise identity ranging from 85% to 90%. In contrast, orphan proteins show lower agreement, with identities between 65% and 75%. Given the high fraction of residues assigned to SSEs, we assessed whether these identity levels could arise from random assignment or instead reflect genuine consensus between methods (see Methods and Appendix A). Even for the least concordant pair of methods (AlphaFold2 and ProtT5, 65% identity), the associated p-value is below 0.05 for 95% of the sequences, confirming that TBM predictions are significantly more similar than expected by chance (Figures S1,S2,S3).

By contrast, marked differences in pLDDT values are observed for residues belonging to SSEs, particularly beta sheets (Figure S5). This observation is consistent with the overall lower pLDDT scores associated with orphan proteins and indicates that, irrespective of their true intrinsic disorder content, TBMs do not model orphan proteins as being more disordered than non-orphan proteins.

## Discussion

Transformer-based models (TBMs) have rapidly become the dominant paradigm for protein structure prediction, largely due to their exceptional performance on proteins embedded in rich evolutionary contexts. In this work, we explicitly tested whether this success extends to genuinely out-of-distribution sequences, focusing on orphan proteins that either originate *de novo* or have diverged beyond detectable homology. Using a curated dataset of Meloidogyne orphan proteins, we show that this regime exposes clear limitations of current TBMs: indeed, tertiary structure predictions are incoherent across different methods, and pLDDT values are significantly lower than for non-orphan proteins. Intrinsic disorder alone does not account for this prediction failure. Whereas there is evidence for *de novo* proteins to be enriched in intrinsic disorder (Wilson et al., 2017), diverged proteins (accounting for 50-80% of orphan proteins) are thought to retain structural similarity with their original protein family. Our results agree with this intuition and show that even for confirmed *de novo* proteins, disorder prediction estimates show an increase only when directly or indirectly calculated with TBMs. Instead, we find that secondary structure is consistently and significantly recovered across models, even when global folds disagree. A potential concern is that orphan proteins are substantially shorter than the typical protein used to benchmark structure predictors, raising the possibility that our observations might be driven by sequence length rather than orphan status per se. We explicitly controlled for this effect by repeating all analyses on a length-matched subset of non-orphan proteins, selected to reproduce the orphan length distribution (Figures S6,S Across all metrics, the results were unchanged. This demonstrates that the observed deficiencies in tertiary structure prediction are not a trivial consequence of shorter sequences, but instead reflect a failure to generalize to sequences lacking identifiable evolutionary context.

Although our dataset is restricted to Meloidogyne orphan proteins, multiple lines of evidence suggest that the conclusions extend more broadly. Comparative studies have shown that many statistical and compositional features of *de novo* proteins are remarkably conserved across taxa, despite their independent evolutionary origins (Basile et al., 2017). In line with this, a recent study on Drosophila *de novo* proteins reports strikingly similar results: low confidence and poor agreement in tertiary structure prediction, and limited explanatory power of intrinsic disorder (Middendorf and Eicholt, 2024). While broader validation across additional clades will be necessary, the convergence of these findings strongly suggests that the observed behavior reflects a general property of TBMs confronted with truly novel protein sequences rather than lineage-specific peculiarities.

These results naturally raise the question of whether the observed failures are inherent to transformer architectures themselves. Recent work suggests that TBMs struggle to learn complex evolutionary and functional abstractions beyond what is directly supported by training data, particularly when evolutionary depth is shallow or absent (Hassan et al., 2025). Complementarily, theoretical and empirical analyses of transformers indicate that they predominantly capture local and mid-range interactions, such as motifs and short structural patterns, rather than encoding truly global context (Zhang et al., 2024), and that they incorporate structural prediction biases (Sarti and Cazals, 2026) (Pratt et al., 2025). This perspective offers a parsimonious explanation for our observations: secondary structure elements, which are largely determined by local sequence patterns, remain accessible to TBMs even in orphan proteins, whereas tertiary structure, requiring long-range coordination and global constraints, cannot be reliably inferred when evolutionary signals are missing.

Taken together, our results suggest a clear conceptual distinction between what TBMs interpolate and what they genuinely generalize. While these models excel at reconstructing folds represented in their training distributions, their predictions for orphan proteins reveal an intrinsic reliance on evolutionary redundancy. Importantly, the partial success observed at the level of secondary structure indicates that TBMs do capture meaningful biophysical regularities, but that these are insufficient to resolve full three-dimensional organization in the absence of homologous constraints. Orphan proteins therefore constitute a stringent and biologically relevant benchmark for probing the limits of modern protein language models, and highlight the need for future architectures or training strategies that better integrate physical principles and global structural reasoning.

## Funding

This research was supported by the joint INRAE-Inria PhD program, which funds the PhD thesis of Er.S.

We are grateful to the Genotoul bioinformatics platform Toulouse Occitanie (Bioinfo Genotoul, DOI: 10.15454/1.5572369328961167E12) as well as to the bioinformatics and genomics platform, BIG, Sophia Antipolis (ISC PlantBIOs, DOI: 10.15454/qyey-ar89) for computing and storage resources.

## Data availability

The original orphan dataset can be found at The dataset including all our analysis has been formatted in the same fashion and is available at All scripts used for the analyses are available at https://doi.org/10.5281/zenodo.18788931

## Author contributions statement

Ed.S. conceived the idea, Er.S. and Ed.S. conducted the analyses, E.D. and D.C. provided the biological expertise and guided the research, Ed.S., Er.S., E.D. and D.C. wrote and reviewed the manuscript.

## Acknowledgments

The authors thank Frédéric Cazals, Mathilde Carpentier, Gianni Liti, Alessandra Carbone and Vincent Mallet for the insightful discussions.

