## Supporting Data and Figures for "Evaluating transformer-based models for structural characterization of orphan proteins"

### Appendix A - p-value calculation of consensual secondary structure assignment

Given two different amounts of predicted alpha-helical, beta-sheet, or coil amino acids, we want to calculate how likely it is for the two different arrangements to agree on a certain number (or percentage) of residue-wise secondary structure annotations. We explicitly calculate all permutations corresponding to a fixed number of alpha-helix and beta-sheet (but not coil) consensual annotations (respectively $n_H$ and $n_B$, given the total number of helix- and beta-annotated residues in both predictions, and the common length of the sequence (respectively $H_1$, $H_2$, $B_1$, $B_2$, and $L$).

Let us first solve the problem when only one variety is present. Given a sequence of length $L$ and two predictions for the total number of alpha-helix residues $0 \leq H_1, H_2 \leq L$, how many permutations result exactly in $n_H$ alpha-helix residues predicted by consensus? To solve this, we have to account for all permutations for choosing $n_H$ out of $H_1$ and $H_2$ residues, as well as the permutations between said elements. Moreover, we have to account for all permutations the remaining $H_1 - n_H$ and $H_2 - n_H$ alpha-helix residues will be paired to coil residues, their internal permutations, and the permutations of the remaining coil residues. This brings us to the following formula:

$$\begin{aligned}\#(n_H|H_1,H_2,L) =& \binom{H_1}{n_H}\binom{H_2}{n_H}\binom{L-H_2}{H_1-n_H}\binom{L-H_1}{H_2-n_H} \\ &\cdot n_H!(H_1-n_H)!(H_2-n_H)!(L-H_1-H_2+n_H)!\end{aligned} \tag{1}$$

Now let's address the complete case with two different moieties $H$ and $B$. The added difficulty is accounting for cross-assignments $H$-$B$ (here $n_{HB}$ if the residue is assigned $H$ in the first and $B$ in the second, and $n_{BH}$ otherwise), which in turn depends on the assignment of $B$ moieties. We choose a recursive approach that explicitly uses the above single-moiety formula. The resulting formula is:

$$\begin{aligned}\#(n_H,n_B|H_1,H_2,B_1,B_2,L) =& \binom{H_1}{n_H}\binom{H_2}{n_H}\binom{B_1}{n_B}\binom{B_2}{n_B} n_H! n_B! \\ &\cdot \sum_{n_{HB}=\ell_{HB}}^{\mathscr{L}_{HB}} \sum_{n_{BH}=\ell_{BH}}^{\mathscr{L}_{BH}} n_{HB}! n_{BH}! (H_1-n_H-n_{HB})!(H_2-n_H-n_{BH})! \\ &\cdot \binom{H_1-n_H}{n_{HB}}\binom{H_2-n_H}{n_{BH}}\binom{B_2-n_B}{n_{HB}}\binom{B_1-n_B}{n_{BH}} \\ &\cdot \#(0|H_1=B_1-n_B-n_{BH}, H_2=B_2-n_B-n_{HB}, L=\mathscr{M})\end{aligned} \tag{2}$$

where

$$\begin{aligned}\ell_{HB} &= \max(0,(H_1-n_H)-(L-H_2-B_2))) \\ \mathscr{L}_{HB} &= \min(H_1-n_H, B_2-n_B) \\ \ell_{BH} &= \max(0,(H_2-n_H)-(L-H_1-B_1))) \\ \mathscr{L}_{BH} &= \min(H_2-n_H, B_1-n_B) \\ \mathscr{M} &= \max(0, L-(H_1+H_2-2n_H)-n_H-n_B)\end{aligned}$$

### Consensual secondary structure assignment

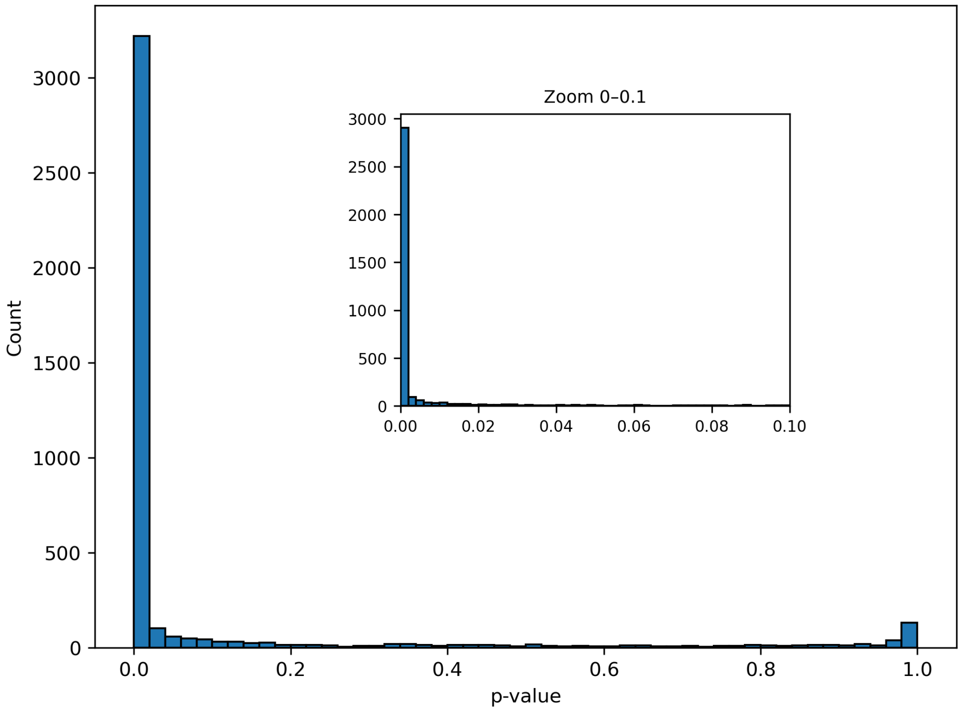


**Figure S1: Significance of consensual secondary structure predictions between AlphaFold2 and ESMFold.** The histogram reports the p-value of the consensus in the secondary structure prediction of each clade 1 orphan protein. The vast majority of cases falls below 0.001.

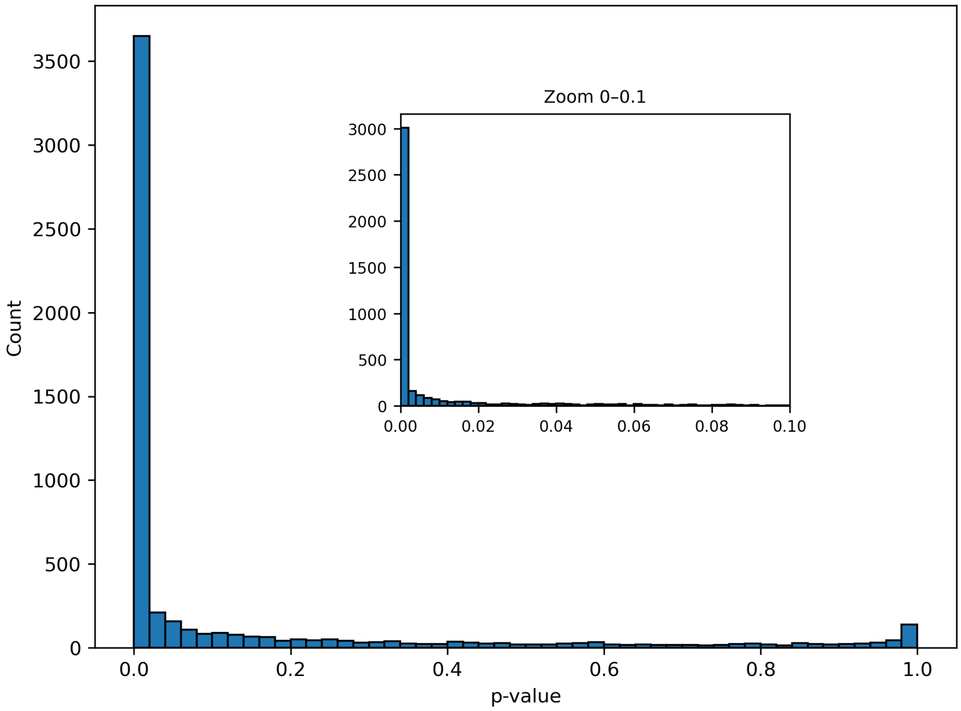


**Figure S2: Significance of consensual secondary structure predictions between ProtT5 and AlphaFold2.** The histogram reports the p-value of the consensus in the secondary structure prediction of each clade 1 orphan protein. The vast majority of cases falls below 0.001.

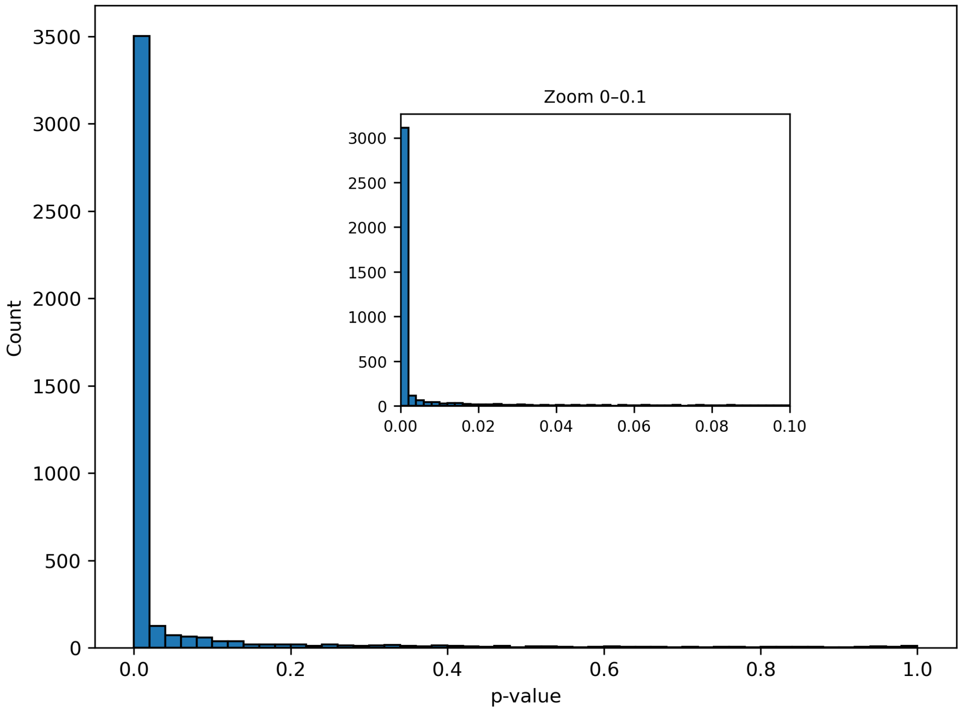


**Figure S3: Significance of consensual secondary structure predictions between ProtT5 and ESMFold**. The histogram reports the p-value of the consensus in the secondary structure prediction of each clade 1 orphan protein. The vast majority of cases falls below 0.001.

### Supplementary figures

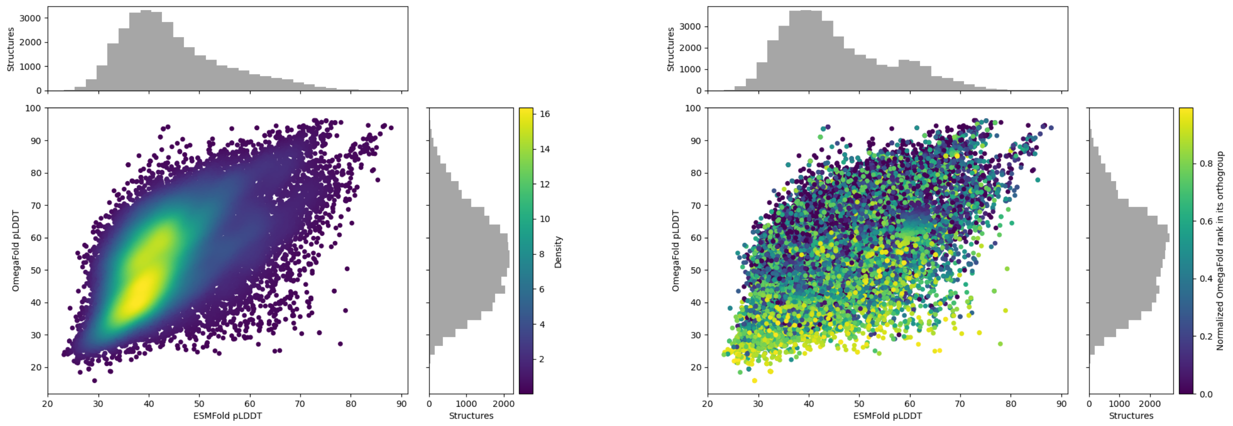


**Figure S4: pLDDT correlation between ESMFold and OmegaFold.** Right: Correlation between the orphan protein structure prediction pLDDT of ESMFold and OmegaFold. Left: structures are colored with respect to their score ranking within their orthogroup, according to OmegaFold.

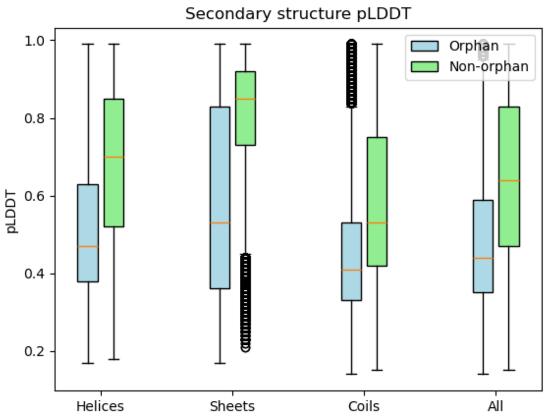


**Figure S5: pLDDT of secondary structure residues for ESMFold. The decrease in pLDDT is consistent throughout the presence or absence of secondary structure.**

#### Sequence length-restricted orphan protein dataset

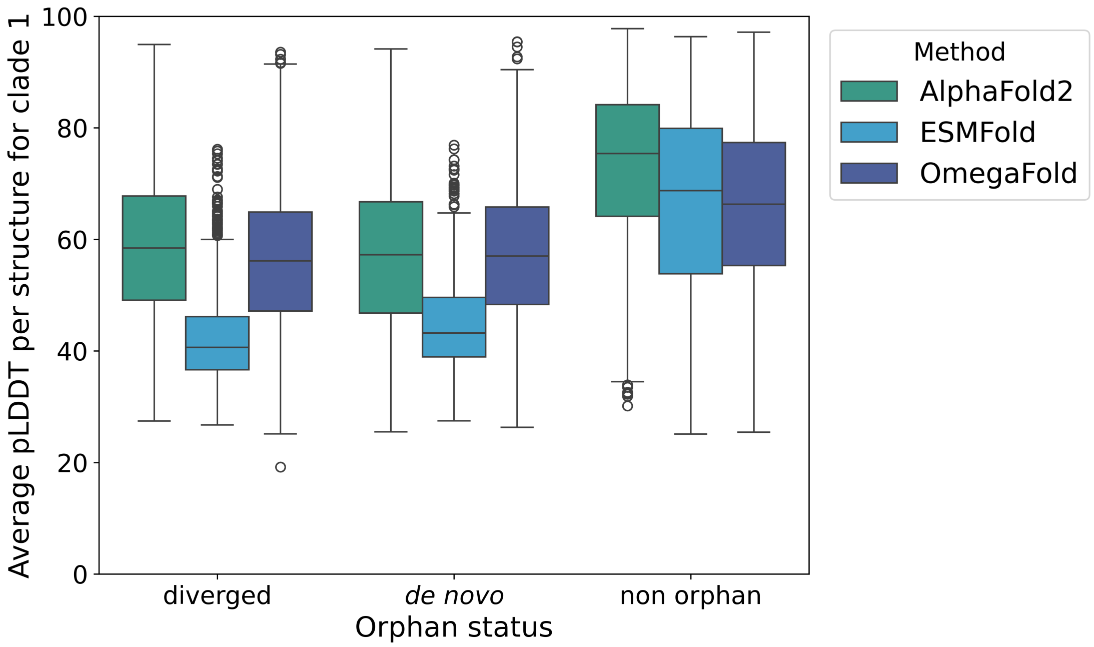


**Figure S6: Average pLDDT of orphan protein structure predictions for three TBMs on same-sequence-length datasets.** Average pLDDT of AlphaFold2, ESMFold and OmegaFold on confirmed diverged and *de novo* proteins. Average pLDDT of a selected same-size dataset of non-orphan proteins with matching sequence lengths. Sequence length is not a relevant factor for our observations.

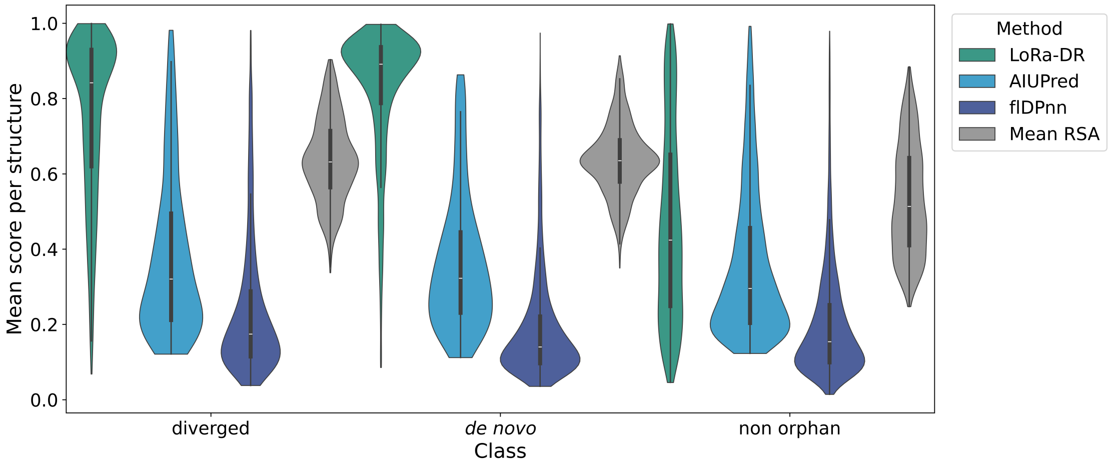


**Figure S7: Predicted intrinsic disorder for orphan and non-orphan proteins on same-sequence-length datasets.** Predicted intrinsic disorder with LoRA-DR, AIUPRed, flDPnn and AlphaFold2 (through RSA calculation). Sequence length does not play a significant role in this analysis.
